# Improved Split Fluorescent Proteins for Endogenous Protein Labeling

**DOI:** 10.1101/137059

**Authors:** Siyu Feng, Sayaka Sekine, Veronica Pessino, Han Li, Manuel D. Leonetti, Bo Huang

## Abstract

Self-complementing split fluorescent proteins (FPs) have been widely used for protein labeling, visualization of subcellular protein localization, and detection of cell-cell contact. To expand this toolset, we have developed a screening strategy for the direct engineering of self-complementing split FPs. Via this strategy, we have generated a yellow-green split-mNeonGreen2_1-10/11_ that improves the ratio of complemented signal to the background of FP_1-10-_expressing cells compared to the commonly used split-GFP_1-10/11,_ as well as a 10-fold brighter red-colored split-sfCherry2_1-10/11_. Based on split-sfCherry2, we have engineered a photoactivatable variant that enables single-molecule localization-based super-resolution microscopy. We have demonstrated dual-color endogenous protein tagging with sfCherry2_11_ and GFP_11_, revealing that endoplasmic reticulum translocon complex Sec61B has reduced abundance in certain peripheral tubules. These new split FPs not only offer multiple colors for imaging interaction networks of endogenous proteins, but also hold the potential to provide orthogonal handles for biochemical isolation of native protein complexes.

## INTRODUCTION

Self-complementing split fluorescent proteins (FPs) are split FP constructs in which the two fragments can associate by themselves to form a fully functional FP without the assistance of other protein-protein interactions. By fusing one fragment on a target protein and detecting its association with the other fragment, these constructs have demonstrated powerful applications in the visualization of subcellular protein localization^1-3^, quantification of protein aggregation^4^, detection of cytosolic peptide delivery^5,6^, identification of cell contacts and synapses ^7,8^, as well as scaffolding protein assembly^3,9,10^. Recently, they have also enabled the generation of large-scale human cell line libraries with fluorescently tagged endogenous proteins through CRISPR/Cas9-based gene editing^11^.

So far, the most commonly used self-complementing split FP was GFP_1-10D7/11M3 OPT_ (which we referr to as GFP_1-10/11_), engineered from super-folder GFP (sfGFP)^12^. With the splitting point between the 10^th^ and 11^th^ β-strands, the resulting GFP_11_ fragment is a 16-amino-acid (a.a.) short peptide. The corresponding GFP_1-10_ fragment remains almost non-fluorescent until complementation, making GFP_1-10/11_ well suited for protein labeling by fusing GFP_11_ to the target protein and over-expressing GFP_1-10_ in the corresponding subcellular compartments. However, there lacks a second, orthogonal split FP system with comparable complementation performance for multicolor imaging and multiplexed scaffolding of protein assembly. Previously, a sfCherry_1-10/11_ system^3^ was derived from super-folder Cherry, an mCherry variant optimized for folding efficiency^13^. However, its overall fluorescent brightness is substantially weaker than an intact sfCherry fusion, potentially due to its limited complementation efficiency^3^. Although two-color imaging with sfCherry_1-10/11_ and GFP_1-10/11_ has been done using tandem sfCherry_11_ to amplify the sfCherry signal for over-expressed targets, it is still too dim to detect most endogenous proteins.

In this paper, we report a screening strategy for the direct engineering of self-complementing split FPs. Using this strategy, we have generated a yellow-green colored mNeonGreen2_1-10/11_ (mNG2) that has an improved ratio of complemented signal to the background of FP_1-10_-expressing cells as compared to GFP_1-10/11_, as well as a red colored sfCherry2_1-10/11_ that is about 10 times as bright as the original sfCherry_1-10/11_. Further, we have engineered a photoactivatable PAsfCherry2_1-10/11_ for single-molecule switching-based super-resolution microscopy. Using these split FPs, we have demonstrated dual-color endogenous protein tagging, which has revealed the reduced abundance of the endoplasmic reticulum (ER) translocon component Sec61B from certain peripheral ER tubules.

## RESULTS

### Engineering split FPs with the spacer-insertion strategy

Inspired by assays previously used to optimize a protease reporter^9^, we devised a general strategy for the engineering of self-complementing split FPs. Specifically, we inserted a 32 a.a. spacer (DVGGGGSEGGGSGGPGSGGEGSAGGGSAGGGS) between the 10^th^ and 11^th^ β-strands of a fluorescent protein (Fig. 1a). This long spacer hinders the folding of the FP, which results in a fluorescence level much lower than its full length counterpart without the spacer. To improve the fluorescence, we then subjected the spacer-inserted FP to multiple rounds of directed evolution in *E. coli*. In each round, the coding sequence was randomly mutagenized or shuffled. Then, the brightest 1 or 2 colonies from each plate were selected for the next round.

**Figure 1.**
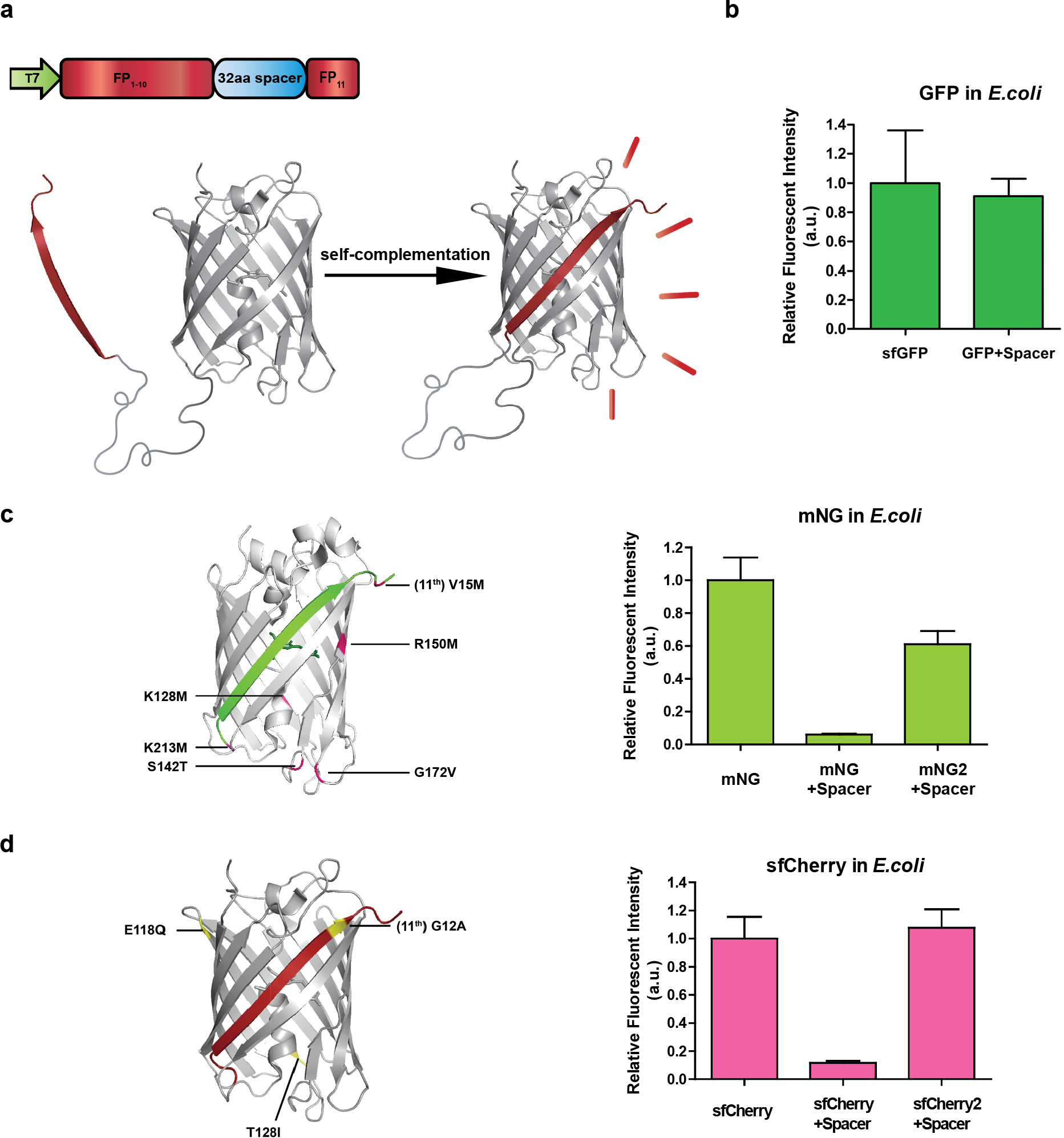
Engineering split FP constructs using the spacer-insertion strategy. **(a)** Construct design and complementation scheme of the spacer-insertion screening system. **(b)** Relative fluorescence intensities of sfGFP and GFP + spacer in *E. coli* colonies. **(c)** Left: schematic diagram of split mNeonGreen2 with 6 mutations highlighted in pink, illustrated on the crystal structure of lanGFP. Right: relative fluorescence intensities of mNG, mNG + spacer, mNG2 + spacer in *E.coli* colonies. **(d)** Left: schematic diagram of split sfCherry2 with 3 mutations highlighted in yellow, illustrated on the crystal structure of sfCherry. Right: relative fluorescence intensities of sfCherry, sfCherry2 + spacer, sfCherry + spacer in *E. coli* colonies. For each bar in all bar graphs, Number of colonies > 400 and error bars are standard deviations.

We first aimed to produce a green-colored split FP that has improved brightness compared to GFP. A recent quantitative assessment of FPs^14^ reported that the brightness of mNeonGreen (mNG)^15^, a yellow-green fluorescent protein derived from *Branchiostoma Lanceolatum*, is more than 2 times higher than sfGFP. mNG also demonstrates good photostablility, acid tolerance and monomeric quality. Guided by the crystal structure of the closely related lanGFP (PDB: 4HVF), we chose to split between 10^th^ and 11^th^ β-strands of mNG and removed the additional GFP-like C terminus (GMDELYK), resulting in a 213-amino-acid fragment which we called mNG_1-10_ and a 16-amino-acid, mNG_11_. Unlike the highly optimized^12^ split GFP_1-10/11_ system, whose fluorescence signal is only slightly reduced with the spacer insertion (Fig. 1b), inserting the spacer between mNG_1-10_ and mNG_11_ drastically reduced its fluorescence signal (Fig 1c). Using our spacer-assisted screening system, after three rounds of random mutagenesis, we identified 5 substitutions in the 1-10 fragment (K128M, S142T, R150M, G172V and K213M) and 1 substitution in 11^th^ strand (V15M) (Fig. 1c). We named this improved mNG, mNG2. In *E.coli* colonies grown on LB-agar plates, spacer-inserted mNG2 demonstrated a 10-fold improvement in brightness after directed evolution, which is ~60% as bright as a full length mNG (Fig. 1c).

To improve the complementation efficiency of split sfCherry, we subjected the spacer-inserted sfCherry to three rounds of random mutagenesis and one round of DNA shuffling. We identified a new variant, named sfCherry2, which contains two mutations on the 1-10 fragment (E118Q and T128I) and one on the 11^th^ strand (G12A) (Fig. 1d). In *E.coli* colonies, spacer-inserted sfCherry2 is ~ 9 times as bright as the spacer-inserted original sfCherry (Fig. 1d). We have also used this strategy to split FusionRed, a red fluorescent protein with minimal cell toxicity and dimerization tendencies^16^. Unfortunately, we have not been able to obtain a brightly fluorescent, spacer-inserted variant even after four rounds of random mutagenesis.

### Protein labeling by mNG2_1-10/11_ in mammalian cells

To test protein labeling using mNG2_11_, we expressed two proteins, histone H2B (H2B) or clathrin light chain A (CLTA) fused to mNG2_11_ in HeLa cells. With the co-expression of mNG2_1-10_, we could correctly image the localization of these proteins, similar to that using GFP_11_ (Fig. 2a). Interestingly, when GFP_1-10_ is expressed by itself, we observed a weak but non-negligible fluorescence background even without the expression of GFP_11_ fragment. Comparing HEK 293T cell lines stably expressing GFP_1-10_ from the strong SFFV promoter and the weak PGK promotor, we found that the background is positively correlated with the GFP_1-10_ expression level (Fig. 2b). This background might be attributed to either weak fluorescence from GFP_1-10_ or elevated cell autofluorescence caused by GFP_1-10_ expression. In contrast, even when using the SFFV promoter, the signal from mNG2_1-10_-expressing cells is indistinguishable from wild-type cell autofluorescence (Fig. 2b).

**Figure 2.**
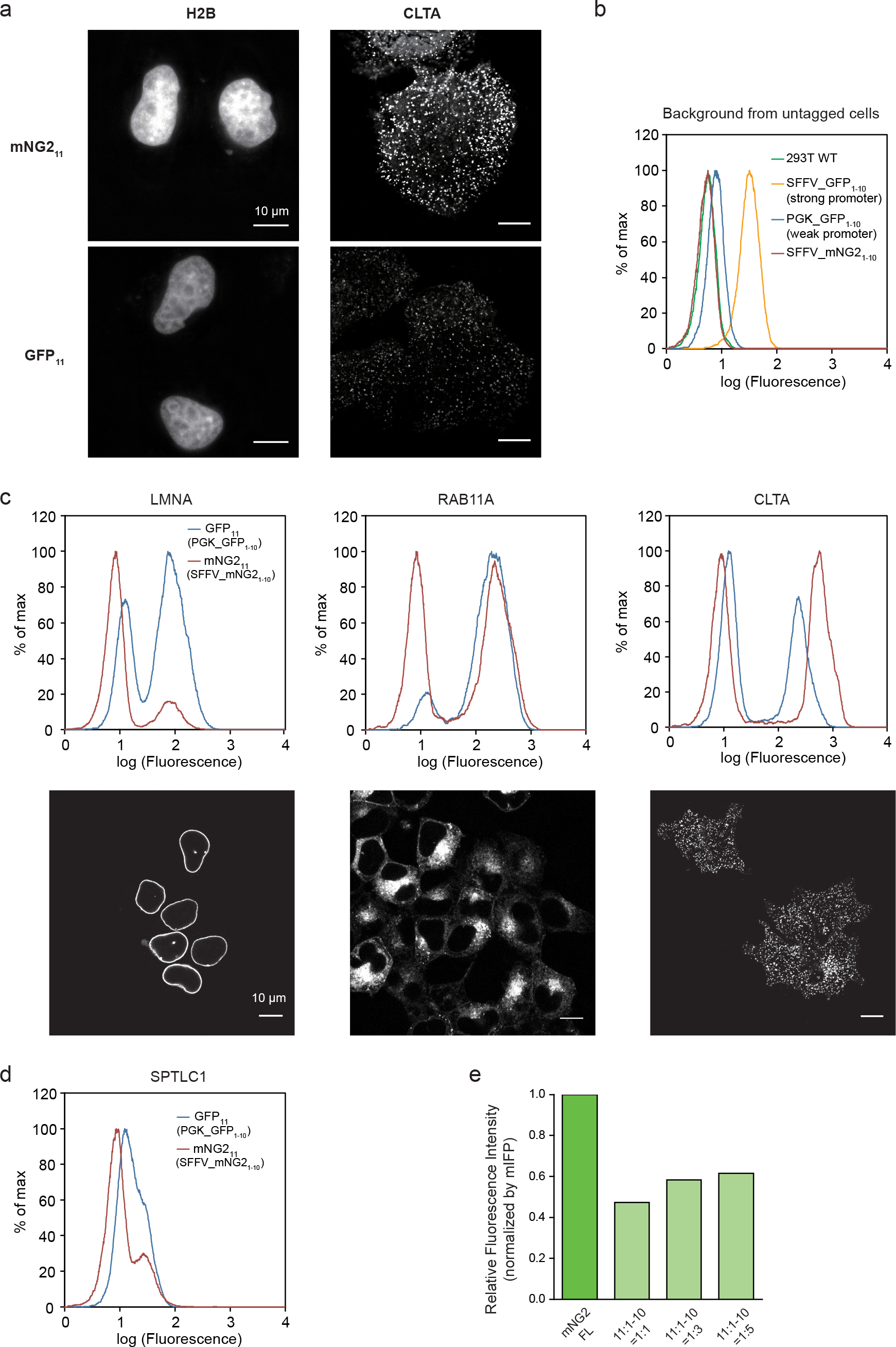
Protein labeling through transient expression or endogenous knock-in using mNG2_11_. **(a)** Fluorescence images of HeLa cells co-expressing either mNG2_11_ or GFP_11_ labeled H2B or CLTA with the corresponding FP_1-10_. Scale bars are 10 μm. **(b)** FACS backgroud from wild type HEK 293T cells, or HEK 293T cells stably expressing GFP_1-10_ (using the stong SFFV promoter or the weak PGK promoter) or mNG2_1-10_ (using SFFV). **(c)** Comparison of tagging endogenous LMNA, RAB11A and CLTA using mNG2_11_ or GFP_11_ knock-in in SFFV-mNG2_1-10_ and PGK-GFP_1-10_ cells, respectively, (flow cytometry histograms) and the corresponding confocal images of mNG2_11_ knock-in cells. Scale bars are 10 μm. **(d)** Comparison of tagging a low-abundance endogenous protein SPTLC1 through mNG2_11_ or GFP_11_ knock-in (flow cytometry histograms). **(e)** Whole cell fluorescence intensity of full length mNG2 and mNG2_11_-CLTA/mNG2_1-10_, measured by flow cytometry and normalized for expression level by mIFP fluorescence signal. Number of cells > 6000.

This low background fluorescence from mNG2_1-10_ is important for the labeling of endogenous proteins using FP_11_ knock-in. Previously, utilizing the small GFP_11_ tag we have developed a scalable scheme to fluorescently tag endogenous proteins in human cell lines by gene editing using electroporation of Cas9/sgRNA ribonucleoprotein (RNP) and a single-stranded donor DNA, followed by enrichment of integrated cells by fluorescence activated cell sorting (FACS)^11^. Similar to GFP_11_, mNG2_11_ is also a 16 a.a. peptide, allowing its DNA and the homology arms to fit in a 200 nucleotide (nt) donor DNA that can be directly obtained from commercial synthesis, which is a key contributor to the efficiency and cost-effectiveness of our method. We compared genetic knock-in using GFP_11_ or mNG2_11_ for three targets: Lamin A/C (LMNA, inner nuclear membrane), RAB11A and CLTA. In all three cases, we observed similar or stronger fluorescence from mNG2_11_ knock-in cells than GFP_11_ knock-in cells. The background from non-integrated cells is substantially lower from mNG2_1-10_, despite the use of the low expression PGK GFP_1-10_ and high expression SFFV mNG2_1-10_. This better separation between FP_11_-integrated and non-integrated cells is especially advantageous for labeling low abundance proteins, for example, SPTLC1 (Serine Palmitoyltransferase Long Chain Base Subunit 1) (Fig. 2d).

Finally, to compare the brightness of mNG2_11_ with that of full length mNG2 on proteins, we constructed plasmids encoding full length mNG2 or mNG2_11_-CLTA fused to mIFP ^17^ through a self-cleaving P2A site, so that expression level differences can be normalized by mIFP signal level. We co-transfected HEK 293T cells with either of the two plasmids and a plasmid expressing mNG2_1-10_. With 11:1-10 transfected DNA ratio increased from 1:1 to 1:3 and 1:5, the normalized whole cell fluorescence by flow cytometry from mNG2_11_ is approximately 50% to 60% of that from full length mNG2 (Fig. 2e). This relative brightness likely indicates the overall effect of complementation efficiency, chromophore maturation and potential purtubation to chromophore environment by the protein split. For reference, we performed the same measurement on GFP_11_ and observed a brightness close to that of the full length sfGFP (Supplementary Fig. 1).

### Protein labeling using sfCherry2_1-10/11_ in mammalian cells

To quantify the improvement of sfCherry2_1-10/11_ over sfCherry_1-10/11_ for mammalian protein labeling and compare their performances with that of full length sfCherry and sfCherry2, we performed a similar measurement as in the previous section. We used TagBFP instead of mIFP for expression level normalization because sfCherry fluorescence bleeds into the mIFP detection channel (Fig. 3a). We found that sfCherry2_1-10/11_ is ~ 10 times as bright as sfCherry_1-10/11_ in mammalian cells. We have also observed that full length sfCherry2 is ~ 50% brighter than sfCherry^13^, suggesting a better overall folding efficiency. The normalized fluorescence signal of sfCherry2_1-10/11_ reached ~ 30% of full length sfCherry and ~ 20% of full length sfCherry2.

**Figure 3.**
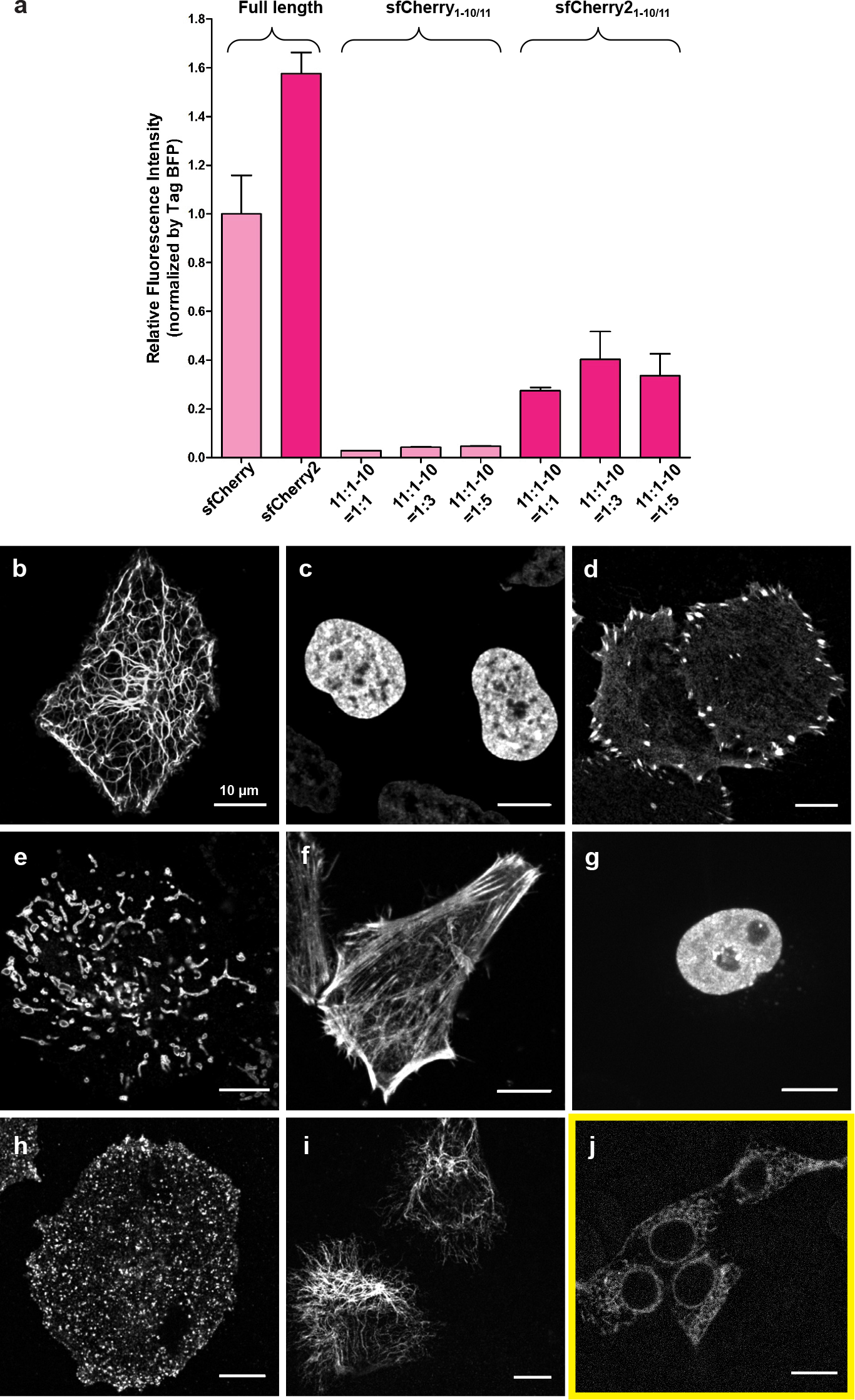
Protein labeling and imaging using sfCherry211. **(a)** Whole cell fluorescence intensity of HEK 293T cells expressing full length sfCherry, full length sfCherry2, sfCherry1-10/11 or sfCherry21-10/11, measured by flow cytometry and normalized for expression level by TagBFP fluorescence signal. Number of cells > 2500. Error bars are s.e.m. **(b-i)** Confocal images of sfCherry211 labeling in HeLa cells co-transfected with sfCherry21-10: keratin (**b**, intermediate filament); histone H2B (**c**, nuclear); zyxin (**d**, focal adhesion); TOMM20 (**e**, mitochondria outer membrane); β-actin (**f**, actin cytoskeleton); heterochromatin protein 1 (**g**, nuclear); clathrin light chain A (**h**, clathrin-coated pits); vimentin (**i**, intermediate filament). **(j)** Confocal image of endogenous Sec61B (ER) labeled by sfCherry211 in HEK 293T cells stably expressing sfCherry21-10. Scale bars are 10 μm.

To test sfCherry2_11_ as a fluorescent tag for live-cell imaging, we constructed mammalian expression vectors encoding target proteins tagged with sfCherry2_11_ at either the N or C terminus and co-expressed each with cytoplasmic sfCherry2_1-10_ in HeLa cells (Figs. 3b-i). For the diverse array of target proteins tested, including nuclear proteins histone H2B and heterochromatin protein 1 (HP1), cytoskeletal proteins β-actin, keratin and vimentin, focal adhesion protein zyxin, CLTA, and mitochondrial outer membrane protein TOMM20, we observed their correct localization. No fluorescent signal was detected with the sfCheryr2_1-10_ fragment alone.

We also demonstrated sfCherry2_11_ labeling of endogenous proteins by knocking it into the ER translocon complex protein Sec61B in HEK 293T cells stably expressing both GFP_1-10_ and sfCherry2_1-10_ (293T^double1-10^). After sorting for red-fluorescence-positive cells by FACS, confocal imaging (Fig. 3j) confirmed the ER specificity of the sfCherry2_11_ signal, which is substantially weaker than the other overexpression cases (Figs. 3b-i). We noticed that many cells also display additional punctate structures, which we have verified to be lysosomes (see the last result section).

This artifact is common for mCherry-derived FPs when expressed for a prolonged period of time, especially when targeting proteins in secretory pathways^18^. It likely results from the fact that mCherry has a beta-barrel structure resisting lysosomal proteolysis and a low pKa, such that it remains fluorescent in the acidic lysosome lumen^19^. We have not observed lysosome labeling in any of the non-ER targets that we have imaged in this work with sfCherry2_11_ labeling.

### Super-resolution microscopy using PAsfCherry2_1-10/11_

With their capability to change their fluorescence properties upon light (usually ultraviolet) irradiation, photoactivatable (PA) FPs ^20^ enable tracking of protein trafficking and imaging of labeled proteins using single-molecule switching-based super-resolution microscopy^21^ (more commonly known as stochastic optical reconstruction microscopy, STORM, or photoactivated localization microscopy, PALM). Previously, mCherry has been engineered into a PA FP (PAmCherry1) after introducing 10 amino acid substitutions^22^. Because these substitutions are located on the 1-10 fragment, we merged them with the sequence of sfCherry2_1-10_ (designated as PAsfCherry2_1-10_). When imaging HEK 293T cells co-expressing sfCherry2_11_-H2B and PAsfCherry2_1-10_, we observed little initial fluorescence in red channel and a large fluorescence increase after 405 nm light irradiation (Fig. 4a), confirming the photoactivation property of complemented sfCherry2_11_ and PAsfCherry2_1-10_.

**Figure 4.**
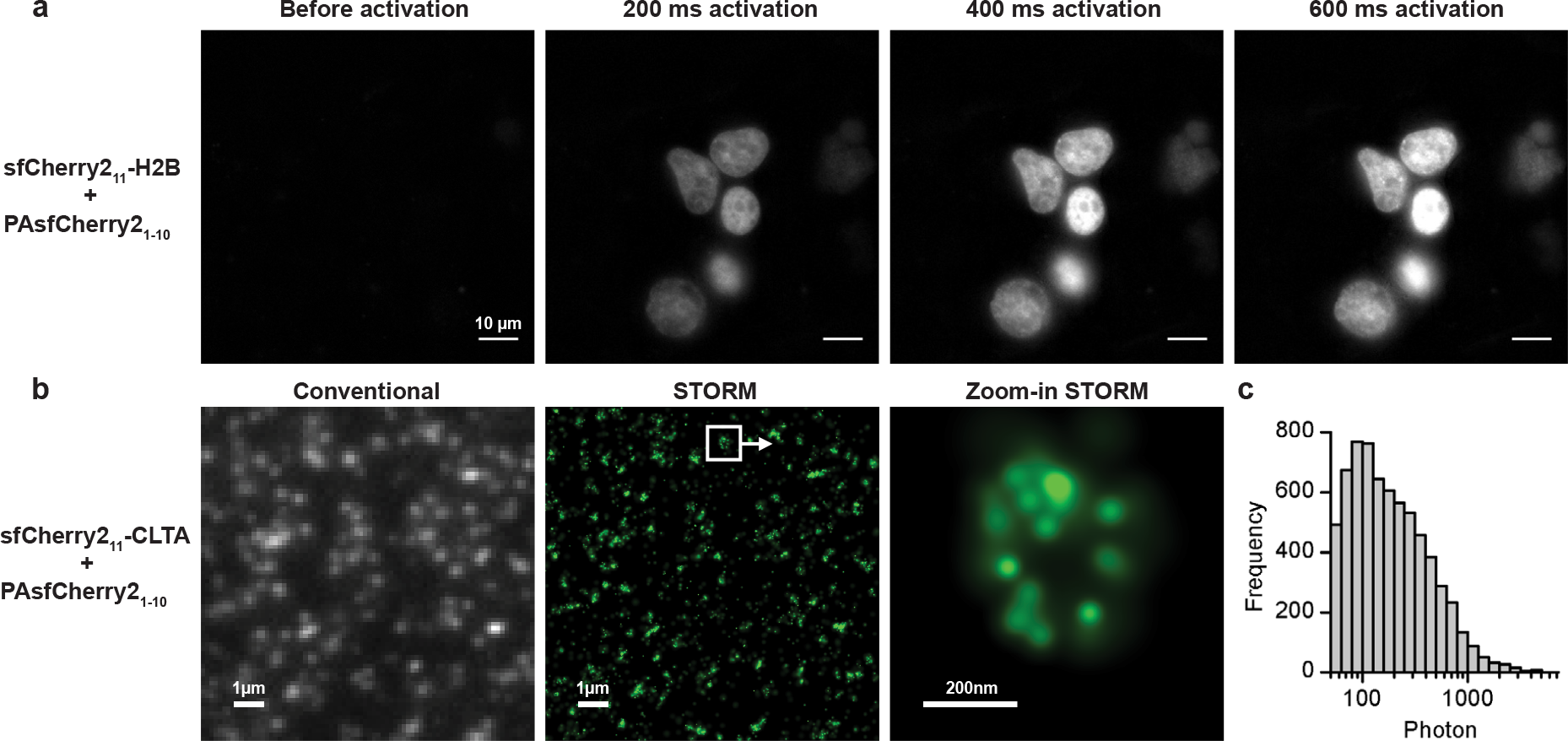
Photoactivatable sfCherry2_1-10/11_ and its use in super-resolution microscopy. **(a)** Photoactivation of H2B labeling in HEK 293T cells co-transfected with sfCherry2_11_-H2B and PAsfCherry2_1-10_. Scale bars: 10 μm. **(b)** Conventional wide-field fluorescence image (left) and super-resolution image (middle/right) of endogenous CLTA. **(c)** Histogram of photon number detected per photoactivation event in **(b)**.

Combining PAsfCherry2_1-10_ with our FP_11_ tag knock-in method, we can easily perform STORM imaging of endogenous proteins tagged by sfCherry_11_, which could avoid potential overexpression artifacts. For demonstration, we imaged endogenous CLTA in HEK 293T cells. Because PAsfCherry2 is non-fluorescent before activation, we knocked-in a tandem GFP_11_-sfCherry2_11_ tag in HEK 293T cells stably expressing GFP_1-10_, so that integrated cells can be isolated using GFP fluorescence signal. We then transfected these cells with PAsfCherry2_1-10_ plasmid for STORM imaging. Compared to conventional wide-field images, clathrin-coated pits can be clearly seen as sub-diffraction-limit objects in the STORM images (Fig. 4b). We were able to collect on average 260 photons in each photoactivation event (Fig. 4c). We note that the labeling of most pits is incomplete, potentially because we labeled only one of the two clathrin light chain genes that have nearly identical functionalities^23^. In addition, non-homozygous knock-in and inefficient complementation between the two fragments can further contribute to the incomplete labeling. Selecting for homozygous knock-in cells and/or the use of tandem sfCherry2_11_ tags^3^ may solve the latter two problems.

### Dual-color endogenous protein tagging in human cells

The orthogonal sfCherry2_11_ now enables two-color imaging of endogenous proteins in order to visualize their differential spatial distribution and interactions. We tested double knock-in of GFP_11_ and sfCherry2_11_ either sequentially or simultaneously in HEK 293T cells stably expressing both GFP_1-10_ and sfCherry2_1-10_ (293T^double1-10^). For sequential knock-in, we first performed electroporation of Cas9 RNP and single-strand donor DNA for GFP_11_, sorted GFP positive cells by FACS, and then knocked-in sfCherry2_11_, followed by a second round of sorting (Fig. 5a). We targeted Sec61B using GFP_11_ and then three proteins with distinctive subcellular localizations using sfCherry2_11_: LMNA, ARL6IP1 (tubular ER), and Sec61B (to verify colocalization). When imaging the double knock-in cells by confocal microscopy, we observed co-labeling of the nuclear envelop by Sec61B-GFP_11_ and LMNA-sfCherry2_11_, almost complete colocalization of Sec61B-GFP_11_ and Sec61B-sfCherry2_11_, and the exclusion of ARL6IP1-sfCherry2_11_ from the nuclear envelope (Fig. 5b).

**Figure 5.**
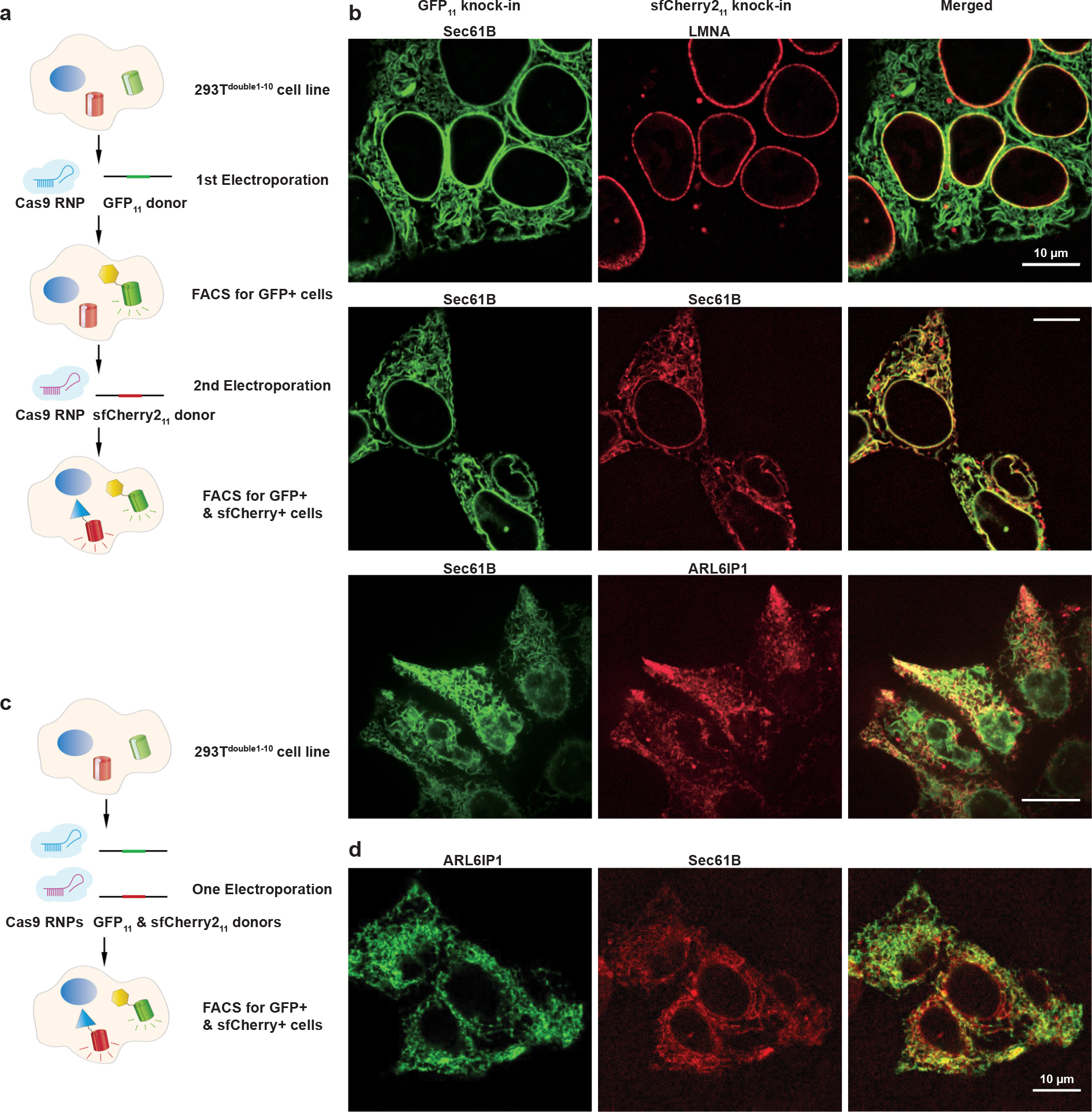
Double labeling and dual-color imaging of endogenous proteins using GFP_11_ and sfCherry2_11_. **(a)** Schemetic diagram for double labeling endogenous proteins through a sequential knock-in strategy. **(b)** Dual-color fluorescence images of GFP_11_/sfCherry2_11_ sequential knocked-in cells. **(c)** Schemetic diagram double labeling endogenous proteins through a simultaneous knock-in strategy. **(d)** Dual-color fluorescence images of GFP_11_/sfCherry2_11_ simultaneous knocked-in cells. Scale bars are 10 µm.

For simultaneous knock-in, we mixed Cas9 RNPs and donor DNAs for both targets in one electroporation reaction and used FACS to enrich cells positive for both green and red signal (Fig. 5c). We chose to tag ARL6IP1 and Sec61B with GFP_11_ and sfCherry2_11_, respectively. We obtained 0.4% double-positive cells, consistent with the multiplication of the 28% and 1.2% efficiency for GFP_11_ and sfCherry2_11_ single knock-in into the ARL6IP1 and Sec61B gene, respectively (Supplementary Fig. 2). The lower efficiency for sfCherry2_11_ knock-in could be attributed to its overall lower fluorescence signal level compared to GFP_11_, making it more difficult to distinguish from the background by FACS. Confocal microscopy of sorted cells showed similar arrangement of the two proteins (Fig. 5d) as in Sec61B-GFP_11_, ARL6IP1-sfCherry2_11_ sequential knock-in.

### Dual-color images reveal peripheral ER with reduced Sec61B

The endoplasmic reticulum (ER) is a large organelle that spreads throughout the cytoplasm as a continuous membrane network of tubules and sheets with a single lumen^24^. The Sec61 complex, which is composed of alpha, beta and gamma subunits, is the central component of the protein translocation apparatus of the ER membrane^25^. Researchers have traditionally used Sec61B as an ER marker for imaging because it is thought to be distributed ubiquitously throughout the ER membrane, including nuclear envelope, sheet-like cisternae, and a polygonal array of tubules^26,27^. However, our dual-color images of endogenous Sec61B and ARL6IP1 in HEK 293T cells using GFP_11_ and sfCherry2_11_, showed that certain peripheral ER tubules marked by ARL6IP1 contain very weak to non-detectable Sec61B signal (Fig. 6a). This large reduction of Sec61B signal in certain ER tubules is clearly visible in cross-section, where the Sec61B signal is at the background level despite Sec61B having the brighter GFP_11_ label than sfCherr2_11_ on ARL6IP1 (Fig. 6b). We have also ruled out the presence of sfCherry2-containing lysosomes in these areas by staining lysosomes with lysotracker (Supplementary Fig. 3). By visual inspection in z maximum projects of 9 confocal images containing a total of 108 cells, we identified that 29 of them contain such peripheral ER tubules lacking strong Sec61B presence. Furthermore, we confirmed this differential distribution of ER membrane proteins by swapping the tags on the two proteins (Fig. 6c).

**Figure 6.**
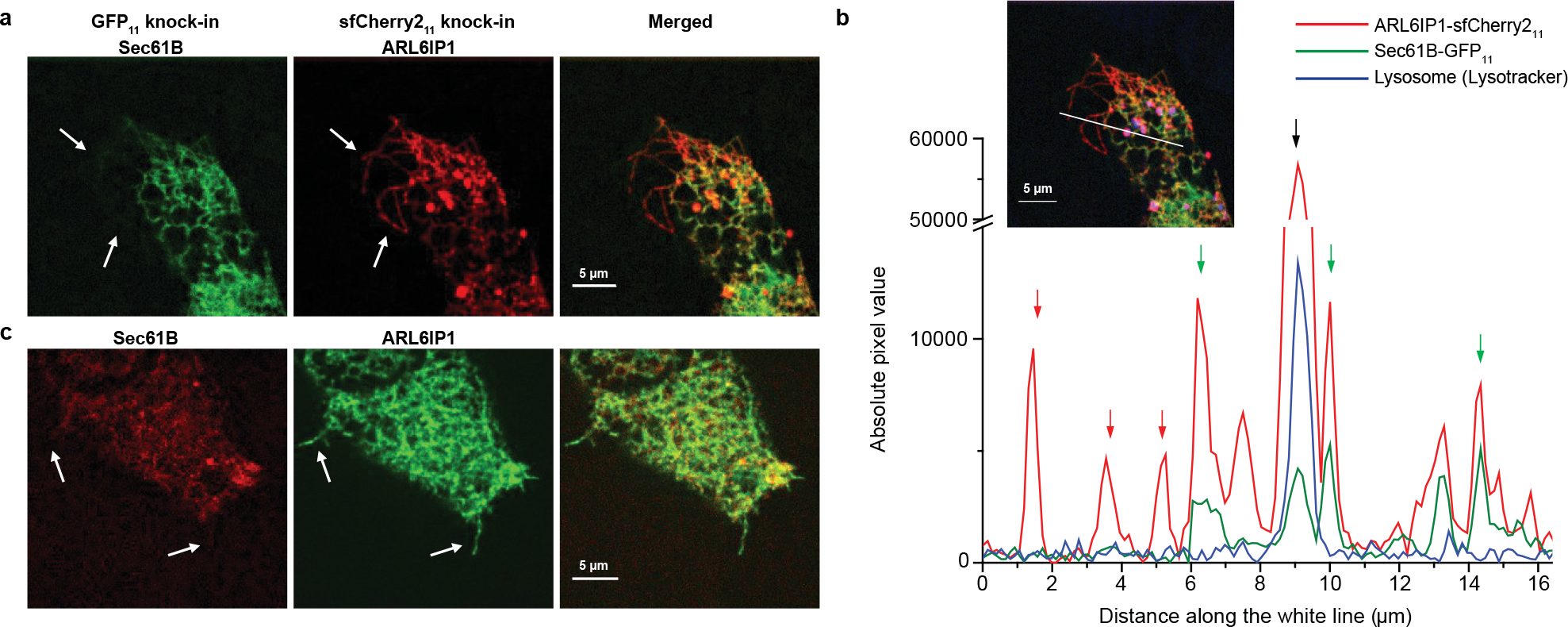
Reduced abundance of Sec61B in certain peripheral ER tubules. **(a)** Dual-color image of a Sec61B-GFP_11_ and ARL6IP1-sfCherry2_11_ knock-in HEK 293T cell. Arrows indicate ER tubules marked by ARL6IP1 but contain much reduced Sec61B signal. **(b)** A cross section in **(a)**, with red arrows indicating petipherial ER tubules with Sec61B signal at the background level, green arrows indicating ER tubules positive for both Sec61B and ARL6IP1, and the black arrow indicating a lysosome marked by LysoTracker Blue. **(c)** Dual-color image of a Sec61B-sfCherry2_11_ and ARL6IP1-GFP_11_ knock-in HEK 263T cell. Scale bars are 5 µm.

## DISCUSSION

In summary, we have devised a simple platform for the engineering of self-complementing split fluorescence proteins. Using this platform, we have developed a bright yellow-green-colored mNG2_1-10/11_ system and substantially increased the performance of the red-colored sfCherry2_1-10/11_. These split constructs have allowed us to obtain two-color or super-resolution images of endogenous proteins and have revealed ER tubules with greatly reduced abundance of the translocon component Sec61B.

Our platform can be easily extended to the engineering of other self-complmenting split FPs with distinct colors (e.g. mTurquoise2, mTagBFP2)^28,29^, good photoactivation performance (e.g. mMaple3, PATagRFP) ^30,31^, or new functionalities (e.g. pH sensitivity)^32^. We note that for non-self-complementing split FPs, which are used to detect protein-protein interactions in bimolecular fluorescence complementation (BiFC) assays^33^, a different engineering platform is needed to ensure minimum affinity between the two FP fragments by themselves. On the other hand, although our optimizated spacer-inserted sfCherry2 has already reached the brightness level of sfCherry without spacer, split sfCherry2_1-10/11_ is still much dimmer compared to full length sfCherry. Further improvement could be possible using a longer or more rigid spacer, which adds more spatial hindrince to complemetation, or a spacer containing the self-cleaving P2A site, which better mimicks the actual split system.

With its higher ratio of complemented signal to the background of FP_1-10_-expressing cells compared to GFP_1-10/11_, our mNG2_1-10/11_ system will be advantageous for tagging low-abundance endogenous proteins^3^. More importantly, it provides an orthogonal handle for scaffolding protein oligomerization^3^ and biochemical isolation of native protein complexes^11^. For the sfCherry2_1-10/11_ system, the fact that we have only observed lysosome puncta when labeling ER proteins suggests that this problem could be potentially resolved by increasing its pKa with rational designs^32^. Moreover, our engineering platform can be easily adapted to generating other red split FPs based on novel bright FPs such as TagRFP-T ^34^, mRuby3 ^35^ or mScarlet^36^.

Given the wide use of Sec61B as a marker to label the entire ER, it is surprising that Sec61B has a substantially reduced abundance in certain peripheral tubular structures labeled by ARL6IP1. It is possible that these tubules, often containing a closed end pointing towards the edge of the cell, serve distinct functions from other ER tubules. Further analysis of their protein compositions and contacts with plasma membrane, cytoskeleton and other organelles may help clarify their functional roles. This observation also suggests that the base of these ER tubules may contain a diffusion barrier that hinders the crossing of Sec61B. Because over-expressing exogenous ER shaping proteins can drastically alter ER morphology^37^, this finding highlights the opportunity of visualizing endogenous proteins to study native interaction networks.

## METHODS

### Mutagenesis and screening

The amino-acid sequence of mNG was obtained from the published literature^15^. Because the crystal structure of mNG has not been determined, we used the lanGFP (PDB: 4HVF) structure as a guide to decide splitting between K213 and T214 at the middle of the loop between 10^th^ and 11^th^ beta-strands. And we also removed the additional 7-residue GFP-like C terminal (GMDELYK) to minimize the size of mNG11 tag. The amino-acid sequence of sfCherry and the split site were from our previous published literature^3^. The spacer-inserted split mNG and split sfCherry were subjected to random mutagenesis using a GeneMorph II Random Mutagenesis Kit (Agilent). The cDNA library pool was transformed into *E.coli* BL21 (DE3) electrocompetent cells (Lucigen) by electroporation using the Gene Pulser Xcell™ Electroporation Systems (BioRad). The expression library was plated on nitrocellulose membrane (Whatman, 0.45um pore size), which was sitting on an LB-agar plate with 30 mg/ml kanamycin. After overnight growth at 37°C, the nitrocellulose membrane was transferred to a new LB-agar plate containing 1mM Isopropyl-β-D-thiogalactoside (IPTG) and 30 mg/ml kanamycin and cultured for 3-6 hours at 37°C to induce the protein expression. Clone screening was performed by imaging the second LB-agar plate using a BioSpectrum Imaging System (UVP). The brightest candidates in each library were pooled (typically 20 to 30 selected from approximately 10,000 colonies) and used as templates for the next round of evolution. For DNA shuffling, we used the method described by Yu et al^38^. Specifically, we digested plasmids containing spacer-inserted sfCherry_1-10/11_ variants with BamHI and XhoI. Fragments of ~ 800bp were purified from 1% agarose gels using zymoclean gel DNA gel recovery kit (Zymo Research). The DNA concentrations were measured and the fragments were mixed at equal amounts for a total of ~2 µg. The mixture was then digested with 0.5 unit DNase I (New England Biolabs) for 13min and terminated by heating at 95°C for 10 min. The DNase I digests were run on a 2% agarose gel, and the size of 50-100 bp fragments were cut out and purified. 10 µl of purified fragments was added to 10 µl of Phusion High-Fidelity PCR Master Mix and reassembled with a PCR program of 30 cycles, with each cycle consisting of 95°C for 60s, 50°C for 60s and 72°C for 30s. After gene reassembly, 1 µl of this reaction was amplified by PCR. The shuffled library was expressed and screened as described above. After the directed evolution was saturated, the brightest clone was selected and the DNA sequences of the constructs were confirmed by sequencing (Quintara Biosciences).

### Molecular cloning

The screening construct was expressed from a pET28a bacterial expression vector and contained a 32-residue spacer (DVGGGGSEGGGSGGPGSGGEGSAGGGSAGGGS) between 10^th^ and 11^th^ beta-strands of FPs. The DNA sequences of mNG1-10_32aaspacer_mNG11 and sfCherry1-10_32aaspacer_sfCherry11 were directly synthesized (Integrated DNA Technologies, IDT). For the nucleotide sequence of identified final mutants of mNG2_1-10_/mNG2_11_ and sfCherry2_1-10_/sfCherry2_11_, see Supplementary Table 1.

The DNAs of mNG2_11_ and sfCherry2_11_ were PCR amplified from identified pET28a constructs (final mutants) using Phusion High-Fidelity DNA Polymerase (Thermo Scientific). The DNAs of histone H2B, clathrin light chain A, keratin, β-actin, zyxin, heterochromatin protein 1, TOMM20, vimentin, laminB, mIFP were subcloned from mEmerald, sfGFP or mIFP fusion plasmids (cDNA source: the Michael Davidson Fluorescent Protein Collection at the UCSF Nikon Imaging Center). The P2A sequence used for all mIFP constructs are GCTACTAACTTCAGCCTGCTGAAGCAGGCTGGAGACGTGGAGGAGAACCCTGGACCT. We performed the following restriction enzyme digestion (amino-acid linker length shown in parentheses for each): histone H2B (10 a.a.): sfGFP sequence between AgeI and BglII (sfGFP-H2B-C-10); clathrin light chain A (15 a.a.): mEmerald sequence between AgeI and BglII (mEmerald-Clathrin-15); keratin (18 a.a): mEmerald sequence between BamHI and NotI (mEmerald-Keratin14-N-18); b-actin (18 a.a.): mEmerald sequence between AgeI and BglII (mEmerald-Actin-C-18); zyxin (6 a.a.): sfGFP sequence between BamHI and NotI (sfGFP-Zyxin-6); TOMM20 (20 a.a.): mEmerald sequence between BamHI and NotI (mEmerald-TOMM20-N-10); vimentin (9 a.a.): mEmerald sequence between BamHI and NotI (mEmerald-Vimentin-N-18); laminB (10 a.a.): mEmerald sequence between AgeI and BglII (mEmerald-LaminB1-10). PCR amplified mNG2_11_ and sfCherry2_11_ were then ligated with the digested vectors using In-Fusion HD Cloning kit (Clontech). For the mammalian expression and lentiviral production, DNAs of mNG2_1-10_ or sfCherry2_1-10_ were directly PCR amplified from identified pET28a constructs (final mutants) and cloned into pcDNA3.1 vectors (HindIII/BamHI) as well as lentiviral pHR-SFFV vector (BamHI/NotI).

To generate the split PAsfCherry2, we introduced 10 photoactivation point mutations (based on PAmCherry1 sequence from literature^22^) to the sfCherry2_1-10_ sequence and cloned the synthesized the PAsfCherry2_1-10_ fragment into a pcDNA3.1 vector. Those substitutions are: E26V/A58T/K69N/L84F/N99K/S148L/I165V/Q167P/L169V/I203R. For the complete nucleotide sequence of PAsfCherry2_1-10_, see Supplementary Table 1.

### Cell culture and generation of stable cell lines

Human HEK 293T and human HeLa cells (obtained from UCSF Cell Culture Facility) were maintained in Dulbecco’s modified Eagle medium (DMEM) with high glucose (UCSF Cell Culture Facility), supplemented with 10% (vol/vol) FBS and 100 µg/ml penicillin/streptomycin (UCSF Cell Culture Facility). All cells were grown at 37°C and 5% CO_2_ in a humidified incubator. Both cell lines have been tested to be free of mycoplasma contamination. For the lentiviral production, HEK 293T cells were plated into 6-well plates one day prior to transfection. 110 ng of pMD2.G plasmid, 890 ng of pCMV-dR8.91 plasmid and 1000 ng of the lentiviral plasmid (pSFFV-mNG2_1-10_, pSFFV-GFP_1-10_ or pSFFV-sfCherry2_1-10_) were cotransfected into HEK 293T in each well using FuGENE HD (Promega) following the manufacturer’s recommended protocol. Lentivirus was harvested 48 h after transfection and were stored immediately in -80°C freezer for future use. To generate the HEK 293T cells stably expressing 1) mNG2_1-10_ or 2) both GFP_1-10_ and sfCherry2_1-10_ (293T^double1-10^), HEK 293T cells were infected with 1) pSFFV-mNG2_1-10_ or 2) both pSFFV-GFP_1-10_ and pSFFV-sfCherry2_1-10_ lentiviruses. 48 hours after infection, we transiently transfected them with 1) mNG2_11_-H2B plasmid or 2) GFP_11_-H2B and sfCherry2_11_-H2B plasmids in order to isolate cells with successful lentiviral integration of 1) mNG2_1-10_ or 2) GFP_1-10_ and sfCherry2_1-10_ by FACS.

### Sample preparation and data analysis in flow cytometry

To characterize the complementation efficiency of split mNG2 and split GFP, we made pSFFV-mIFP_P2A_FP[full length] and pSFFV-mIFP_P2A_FP_11_-CLTA constructs. Corresponding to each bar in Fig. 2e and Supplementary Fig. 1a, HEK 293T cells grown on 48-well plate (Eppendorf) were cotransfected with 1) 80ng pSFFV-mIFP_P2A_mNG2[full length] with 80ng pSFFV-mNG2_1-10_, 2) 80ng pSFFV-mIFP_P2A_mNG2_11_-CLTA with 80ng pSFFV-mNG2_1-10_, 3) 80ng pSFFV-mIFP_P2A_mNG2_11_-CLTA with 160ng pSFFV-mNG2_1-10_, 4) 80ng pSFFV-mIFP_P2A_mNG2_11_-CLTA with 240ng pSFFV-mNG2_1-10_. 5) 80ng pSFFV-mIFP_P2A_sfGFP[full length] with 80ng pSFFV-GFP_1-10_, 6) 80ng pSFFV-mIFP_P2A_GFP_11_-CLTA with 80ng pSFFV-GFP_1-10_, 7) 80ng pSFFV-mIFP_P2A_GFP_11_-CLTA with 160ng pSFFV-GFP_1-10_, 8) 80ng pSFFV-mIFP_P2A_GFP_11_-CLTA with 240ng pSFFV-GFP_1-10_. We varied the ratio of FP_11_ fragment to FP_1-10_ fragment from 1:1 to 1:3 to 1:5, to verify the saturation of complementation.

To characterize the increased fluorescence intensity of split sfCherry2 versus original split sfCherry, we made pSFFV-FP_TagBFP and pSFFV-FP_11__TagBFP constructs. Corresponding to each bars in Fig. 3a, HEK 293T cells grown on 24-well plate (Eppendorf) were transfected using with 1) 500ng pSFFV-sfCherry_TagBFP, 2) 500ng pSFFV-sfCherry2_TagBFP, 3) 500ng pSFFV-sfCherry_11__TagBFP with 500ng pSFFV-sfCherry_1-10_, 4) 500ng pSFFV-sfCherry_11__TagBFP with 1000ng pSFFV-sfCherry_1-10_, 5) 500ng pSFFV-sfCherry_11__TagBFP with 1500ng pSFFV-sfCherry_1-10_, 6) 500ng pSFFV-sfCherry2_11__TagBFP with 500ng pSFFV-sfCherry2_1-10_, 7) 500ng pSFFV-sfCherry2_11__TagBFP with 1000ng pSFFV-sfCherry2_1-10_, 8) 500ng pSFFV-sfCherry2_11__TagBFP with 1500ng pSFFV-sfCherry2_1-10_.

Analytical flow cytometry was carried out on a LSR II instrument (BD Biosciences) and cell sorting on a FACSAria II (BD Biosciences) in Laboratory for Cell Analysis at UCSF. Flow cytometry data analysis (gating and normaliztion) was done using the FlowJo software (FlowJo LLC) and plotted in GraphPad Prism.

### Fluorescence Microscopy

We transfected human HeLa cells grown on an 8-well glass bottom chamber (Thermo Fisher Scientific) using FuGene HD (Promega). In order to achieve better cell attachment, 8-well chamber was coated with Fibronectin (Sigma-Aldrich) for one hour before seeding cells. Total plasmid amount of 120ng per well with the FP_11_ to FP_1-10_ ratio in 1:2 was used to achieve optimal labeling. Thirty-six to forty-eight hours after transfection, live cells were imaged and then fixed with 4% paraformaldehyde. For lysosome staining, LysoTracker^®^ Blue DND-22 (Thermo Fisher Scientific) was added directly to the culture medium (50 nM final concentration) and incubate for 30 min before imaging.

Most of live-cell imaging was acquired on an inverted Nikon Ti-E microscope (UCSF Nikon Imaging Center), a Yokogawa CSU-W1 confocal scanner unit, a Plan Apo VC 100x/1.4NA oil immersion objective, a stage incubator, an Andor Zyla 4.2 sCMOS or an Andor iXon Ultra DU888 EM-CCD camera and MicroManager software. PAsfCherry2 photoactivation in H2B labeling and split mNG2 versus split GFP comparison in H2B labeling were imaged on a Nikon Ti-E inverted wide-field fluorescence microscope equipped with an LED light source (Excelitas X-Cite XLED1), a 100X NA 1.40 PlanApo oil immersion objective, a motorized stage (ASI) and an sCMOS camera (Hamamatsu Flash 4.0). Microscopy images were subjected to background subtraction using a rolling ball radius of 100 pixels in ImageJ Fiji ^39^ software. Analysis of conventional fluorescence microscopy images were performed in ImageJ.

### STORM image acquisition and analysis

Super-resolution images were collected using a TIRF-STORM microscope, home-built from a Nikon Eclipse Ti-E inverted microscope. A 405 nm activation laser (OBIS 405, Coherent), 488 nm imaging laser (OBIS 488, Coherent) and a 561 nm imaging laser (Sapphire 561, Coherent) were aligned, expanded, and focused at the back focal plane of the UPlanSApo 1.4 NA 100x oil immersion objective (Olympus). Images were recorded with an electron multiplying CCD camera (iXon+ DU897E-C20-BV, Andor), and processed with a home-written software. The OBIS lasers were controlled directly by the computer whereas the Sapphire 561 nm laser was shuttered using an acoustic optical modular (Crystal Technology). A quadband dichroic mirror (ZT405/488/561/640rpc, Chroma) and a band-pass filter (ET525/50m, Chroma for 488 nm and ET595/50nm, Chroma for 561 nm) separated the fluorescence emission from the excitation light. Because PAsfCherry2 is initially in a dark state, knock-in cells were carefully located through GFP, using low green intensity (11 μW) and a wide-field setting (5 Hz), so as to minimize potential bleaching of PAsfCherry2. Cells were then subjected to STORM imaging, and if fluorophores appeared in the red channel, it was clear that they were indeed expressing both fluorescent proteins. Maximum laser power used during STORM measured before the objective was 16 μW for 405 nm and 25 mW for 561 nm. By increasing the power on the 405 nm laser (zero to 16 μW) during imaging, PAsfCherry2 was photoconverted from a dark to a red state, and could be observed as single fluorophores. These images were recorded at a frame rate of 30 Hz, with an EMCCD camera gain of 60. During image acquisition, the axial drift of the microscope stage was stabilized by a home-built focus stabilization system utilizing the reflection of an IR laser off the sample. Frames were collected until sample was bleached. Analysis of the STORM images was performed on the Insight3 software ^40^. Cells were imaged in PBS buffer.

### Preparation and electroporation of Cas9/sgRNA RNP

All synthetic nuclei acid reagents were purchased from Integrated DNA Technologies (IDT). sgRNAs and Cas9/sgRNA RNP complexes were prepared using the following procedure^11^. sgRNAs were obtained by *in vitro* transcribing DNA templates containing a T7 promoter (TAATACGACTCACTATAG), the gene-specific 20-nt sgRNA sequence and a common sgRNA scaffold region. DNA templates were generated by overlapping PCR using a set of 4 primers: 3 primers common to all reactions (forward primer T25: 5’-TAA TAC GAC TCA CTA TAG -3’; reverse primer BS7: 5’-AAA AAA AGC ACC GAC TCG GTG C -3’ and reverse primer ML611: 5’-AAA AAA AGC ACC GAC TCG GTG CCA CTT TTT CAA GTT GAT AAC GGA CTA GCC TTA TTT AAA CTT GCT ATG CTG TTT CCA GCA TAG CTC TTA AAC -3’) and one gene-specific primer (forward primer 5’-TAA TAC GAC TCA CTA TAG NNN NNN NNN NNN NNN NNN NNG TTT AAG AGC TAT GCT GGA A -3’). For each template, a 100-μL PCR was performed with iProof High-Fidelity Master Mix (Bio-Rad) reagents with the addition of 1 μM T25, 1μM BS7, 20 nM ML611 and 20 nM gene-specific primer. The PCR product was purified and eluted in 12 μL of RNAse-free DNA buffer (2 mM Tris pH 8.0 in DEPC-treated water). Next, a 100-μL *in vitro* transcription reaction was performed with 300 ng DNA template and 1000 U of T7 RNA polymerase in buffer containing 40 mM Tris pH 7.9, 20 mM MgCl_2_, 5 mM DTT, 2 mM spermidine and 2 mM of each NTP (New England BioLabs). Following a 4h incubation at 37°C, the sgRNA product was purified and eluted in 15 μL of RNAse-free RNA buffer (10 mM Tris pH 7.0 in DEPC-treated water). The sgRNA was quality-checked by running 5 pg of the product on a 10% polyacrylamide gel containing 7 M urea (Novex TBE-Urea gels, ThermoFisher Scientific).

For the knock-in of mNG2_11_, sfCherry2_11_ or GFP_11_, 200-nt homology-directed recombination (HDR) templates were ordered in single-stranded DNA (ssDNA) form as ultramer oligos (IDT). For knock-in of GFP_11_-sfCherry2_11_in tandem, HDR template was ordered in double-stranded (dsDNA) form as gBlock fragments (IDT) and processed to ssDNA as described below. For the complete set of DNA sequence used for sgRNA in vitro transcription or HDR templates, see Supplementary Tables 2 and 3.

Cas9 protein (pMJ915 construct, containing two nuclear localization sequences) was expressed in *E.coli* and purified by the University of California Berkeley Macrolab^41^. 293T mNG2_1-10_ stable cells or 293T^double1-10^ cells were treated with 200 ng/mL nocodazole (Sigma) for ~15 hours before electroporation to increase HDR efficiency^42^. Cas9/sgRNA RNP complexes were assembled with 100 pmol Cas9 protein and 130 pmol sgRNA just prior to electroporation and combined with HDR template in a final volume of 10 μL. Electroporation was carried out in Amaxa 96-well shuttle Nuleofector device (Lonza) using SF-cell line reagents (Lonza). Nocodazole-treated cells were resuspended to 10^4^ cells/μL in SF solution immediately prior to electroporation. For each sample, 20 μL of cells was added to the 10 μL RNP/template mixture. Cells were immediately electroporated using the CM130 program and transferred to 24-well plate with pre-warmed medium. Electroporated cells were cultured for 5-10 days prior to FACS selection of integrated cells.

### Preparation of sfCherry211-GFP11-CLTA ssDNA Template

sfCherry2_11_-GFP_11_-CLTA ssDNA template was prepared from a commercial dsDNA fragment (gBlock, IDT) containing the template sequence preceded by a T7 promoter^11^. The dsDNA fragment was amplified by PCR using Kapa HiFi reagents (Kapa Biosystems) and purified using SPRI beads (AMPure XP resin, Beckman Coulter) at a 1:1 DNA:resin volume ratio (following manufacturer’s instructions) and eluted in 25 μL RNAse-free water. Next, RNA was produced by *in vitro* transcription using T7 HiScribe reagents (New England BioLabs). Following a 4 h reaction at 37°C, the mixture was treated with 4U TURBO DNAse (ThermoFisher Scientific) and incubated for another 15 min at 37°C. The RNA product was then purified using SPRI beads and eluted in 60 μL RNAse-free water. DNA:RNA hybrid was then synthesized by reverse transcription using Maxima H RT reagents (ThermoFisher Scientific). Finally, ssDNA was made by hydrolyzing the RNA strand through the addition of 24 μL NaOH solution (0.5 M NaOH + 0.25 M EDTA, in water) and incubation at 95°C for 10 min. The final ssDNA product was purified using SPRI beads at a 1:1.2 DNA:resin volume ratio and eluted in 15 μL water.

### Data Availability

The data that support the findings of this study are available from the corresponding author upon reasonable request. All relevant DNA sequences are listed in the Supplementary Information.

## ACKNOWLEDGEMENT

We thank Daichi Kamiyama and David A. Brown in the Bo Huang laboratory (UCSF) for advice on molecular cloning and other extensive discussion, Dan Yu in the Xiaokun Shu laboratory (UCSF) for advice on mutagenesis and *E.coli* library screening, and Joseph DeRisi (UCSF) for generously sharing the nucleofector device with us. This work is supported by the National Institutes of Health Director’s New Innovator Award DP2OD008479 (to B.H.) and R21EB022798 (to B.H., S.F. and V.P.), and by the W. M. Keck Foundation Medical Research Grant (to B.H. and S.S.). V.P. is supported by a predoctoral fellowship from the American Heart Association. B.H. is a Chan Zuckerberg Biohub Investigator.

## AUTHOR CONTRIBUTIONS

S.F. and B.H. conceived and designed the research; S.F. performed the screening and protein labeling experiments; S.F., H.L. and M.L. performed the mNG2_11_ knock-in and imaging experiment; S.F. and S.S. performed the dual-color knock-in and imaging experiments; S.F. and V.P. performed the STORM experiments and analysis; S.F. analyzed the data; S.F. and B.H. wrote the manuscript.

## REFERENCES

1 Kaddoum, L., Magdeleine, E., Waldo, G. S., Joly, E. & Cabantous, S. One-step split GFP staining for sensitive protein detection and localization in mammalian cells. Biotechniques 49, 727–736 (2010).

2 Van Engelenburg, S. B. & Palmer, A. E. Imaging type-III secretion reveals dynamics and spatial segregation of Salmonella effectors. Nat Methods 7, 325–330 (2010).

3 Kamiyama, D. et al. Versatile protein tagging in cells with split fluorescent protein. Nat Commun 7, 11046 (2016).

4 Chun, W., Waldo, G. S. & Johnson, G. V. Split GFP complementation assay: a novel approach to quantitatively measure aggregation of tau in situ: effects of GSK3beta activation and caspase 3 cleavage. J Neurochem 103, 2529–2539 (2007).

5 Schmidt, S. et al. Detecting Cytosolic Peptide Delivery with the GFP Complementation Assay in the Low Micromolar Range. Angew Chem Int Ed Engl 54, 15105–15108 (2015).

6 Milech, N. et al. GFP-complementation assay to detect functional CPP and protein delivery into living cells. Sci Rep 5, 18329 (2015).

7 Feinberg, E. H. et al. GFP Reconstitution Across Synaptic Partners (GRASP) defines cell contacts and synapses in living nervous systems. Neuron 57, 353–363 (2008).

8 Macpherson, L. J. et al. Dynamic labelling of neural connections in multiple colours by trans-synaptic fluorescence complementation. Nat Commun 6, 10024 (2015).

9 To, T. L. et al. Rationally designed fluorogenic protease reporter visualizes spatiotemporal dynamics of apoptosis in vivo. Proc Natl Acad Sci U S A 112, 3338–3343 (2015).

10 Kim, Y. E., Kim, Y. N., Kim, J. A., Kim, H. M. & Jung, Y. Green fluorescent protein nanopolygons as monodisperse supramolecular assemblies of functional proteins with defined valency. Nat Commun 6, 7134 (2015).

11 Leonetti, M. D., Sekine, S., Kamiyama, D., Weissman, J. S. & Huang, B. A scalable strategy for high-throughput GFP tagging of endogenous human proteins. Proc Natl Acad Sci U S A 113, E3501–3508 (2016).

12 Cabantous, S., Terwilliger, T. C. & Waldo, G. S. Protein tagging and detection with engineered self-assembling fragments of green fluorescent protein. Nat Biotechnol 23, 102–107 (2005).

13 Nguyen, H. B., Hung, L. W., Yeates, T. O., Terwilliger, T. C. & Waldo, G. S. Split green fluorescent protein as a modular binding partner for protein crystallization. Acta Crystallogr D Biol Crystallogr 69, 2513–2523 (2013).

14 Cranfill, P. J. et al. Quantitative assessment of fluorescent proteins. Nat Methods 13, 557–562 (2016).

15 Shaner, N. C. et al. A bright monomeric green fluorescent protein derived from Branchiostoma lanceolatum. Nat Methods 10, 407–409 (2013).

16 Shemiakina, II et al. A monomeric red fluorescent protein with low cytotoxicity. Nat Commun 3, 1204 (2012).

17 Yu, D. et al. A naturally monomeric infrared fluorescent protein for protein labeling in vivo. Nat Methods 12, 763–765 (2015).

18 Costantini, L. M. et al. A palette of fluorescent proteins optimized for diverse cellular environments. Nat Commun 6, 7670 (2015).

19 Huang, L., Pike, D., Sleat, D. E., Nanda, V. & Lobel, P. Potential pitfalls and solutions for use of fluorescent fusion proteins to study the lysosome. PLoS One 9, e88893 (2014).

20 Lippincott-Schwartz, J. & Patterson, G. H. Photoactivatable fluorescent proteins for diffraction-limited and super-resolution imaging. Trends Cell Biol 19, 555–565 (2009).

21 Huang, B., Babcock, H. & Zhuang, X. Breaking the diffraction barrier: super-resolution imaging of cells. Cell 143, 1047–1058 (2010).

22 Subach, F. V. et al. Photoactivatable mCherry for high-resolution two-color fluorescence microscopy. Nat Methods 6, 153–159 (2009).

23 Brodsky, F. M. Diversity of clathrin function: new tricks for an old protein. Annu Rev Cell Dev Biol 28, 309–336 (2012).

24 English, A. R., Zurek, N. & Voeltz, G. K. Peripheral ER structure and function. Curr Opin Cell Biol 21, 596–602 (2009).

25 Greenfield, J. J. & High, S. The Sec61 complex is located in both the ER and the ER-Golgi intermediate compartment. J Cell Sci 112 (Pt 10), 1477–1486 (1999).

26 Friedman, J. R., Webster, B. M., Mastronarde, D. N., Verhey, K. J. & Voeltz, G. K. ER sliding dynamics and ER-mitochondrial contacts occur on acetylated microtubules. J Cell Biol 190, 363–375 (2010).

27 Nixon-Abell, J. et al. Increased spatiotemporal resolution reveals highly dynamic dense tubular matrices in the peripheral ER. Science 354, aaf3928 (2016).

28 Goedhart, J. et al. Structure-guided evolution of cyan fluorescent proteins towards a quantum yield of 93%. Nat Commun 3, 751 (2012).

29 Subach, O. M., Cranfill, P. J., Davidson, M. W. & Verkhusha, V. V. An enhanced monomeric blue fluorescent protein with the high chemical stability of the chromophore. PLoS One 6, e28674 (2011).

30 Wang, S., Moffitt, J. R., Dempsey, G. T., Xie, X. S. & Zhuang, X. Characterization and development of photoactivatable fluorescent proteins for single-molecule-based superresolution imaging. Proc Natl Acad Sci U S A 111, 8452–8457 (2014).

31 Subach, F. V., Patterson, G. H., Renz, M., Lippincott-Schwartz, J. & Verkhusha, V. V. Bright monomeric photoactivatable red fluorescent protein for two-color super-resolution sptPALM of live cells. J Am Chem Soc 132, 6481–6491 (2010).

32 Shen, Y., Rosendale, M., Campbell, R. E. & Perrais, D. pHuji, a pH-sensitive red fluorescent protein for imaging of exo- and endocytosis. J Cell Biol 207, 419–432 (2014).

33 Hu, C. D., Chinenov, Y. & Kerppola, T. K. Visualization of interactions among bZIP and Rel family proteins in living cells using bimolecular fluorescence complementation. Mol Cell 9, 789–798 (2002).

34 Shaner, N. C. et al. Improving the photostability of bright monomeric orange and red fluorescent proteins. Nat Methods 5, 545–551 (2008).

35 Bajar, B. T. et al. Improving brightness and photostability of green and red fluorescent proteins for live cell imaging and FRET reporting. Sci Rep 6, 20889 (2016).

36 Bindels, D. S. et al. mScarlet: a bright monomeric red fluorescent protein for cellular imaging. Nat Methods 14, 53–56 (2017).

37 Yamamoto, Y., Yoshida, A., Miyazaki, N., Iwasaki, K. & Sakisaka, T. Arl6IP1 has the ability to shape the mammalian ER membrane in a reticulon-like fashion. Biochem J 458, 69–79 (2014).

38 Yu, D. et al. An improved monomeric infrared fluorescent protein for neuronal and tumour brain imaging. Nat Commun 5, 3626 (2014).

39 Schindelin, J. et al. Fiji: an open-source platform for biological-image analysis. Nat Methods 9, 676–682 (2012).

40 Huang, B., Wang, W., Bates, M. & Zhuang, X. Three-dimensional super-resolution imaging by stochastic optical reconstruction microscopy. Science 319, 810–813 (2008).

41 Jinek, M. et al. A programmable dual-RNA-guided DNA endonuclease in adaptive bacterial immunity. Science 337, 816–821 (2012).

42 Lin, S., Staahl, B. T., Alla, R. K. & Doudna, J. A. Enhanced homology-directed human genome engineering by controlled timing of CRISPR/Cas9 delivery. Elife 3, e04766 (2014).

